# Single cell and spatial alternative splicing analysis with long read sequencing

**DOI:** 10.1101/2023.02.23.529769

**Authors:** Yuntian Fu, Heonseok Kim, Jenea I. Adams, Susan M. Grimes, Sijia Huang, Billy T. Lau, Anuja Sathe, Paul Hess, Hanlee P. Ji, Nancy R. Zhang

## Abstract

Long-read sequencing has become a powerful tool for alternative splicing analysis. However, technical and computational challenges have limited our ability to explore alternative splicing at single cell and spatial resolution. The higher sequencing error of long reads, especially high indel rates, have limited the accuracy of cell barcode and unique molecular identifier (UMI) recovery. Read truncation and mapping errors, the latter exacerbated by the higher sequencing error rates, can cause the false detection of spurious new isoforms. Downstream, there is yet no rigorous statistical framework to quantify splicing variation within and between cells/spots. In light of these challenges, we developed Longcell, a statistical framework and computational pipeline for accurate isoform quantification for single cell and spatial spot barcoded long read sequencing data. Longcell performs computationally efficient cell/spot barcode extraction, UMI recovery, and UMI-based truncation- and mapping-error correction. Through a statistical model that accounts for varying read coverage across cells/spots, Longcell rigorously quantifies the level of inter-cell/spot versus intra-cell/ spot diversity in exon-usage and detects changes in splicing distributions between cell populations. Applying Longcell to single cell long-read data from multiple contexts, we found that intra-cell splicing heterogeneity, where multiple isoforms co-exist within the same cell, is ubiquitous for highly expressed genes. On matched single cell and Visium long read sequencing for a tissue of colorectal cancer metastasis to the liver, Longcell found concordant signals between the two data modalities. Finally, on a perturbation experiment for 9 splicing factors, Longcell identified regulatory targets that are validated by targeted sequencing.

## Introduction

Alternative splicing is a pervasive gene regulatory mechanism which affects more than 90% of multi-exon human genes. It is a key regulatory step in gene expression that allows a limited genome to express an impressive diversity of coding and non-coding RNAs^1^. Alternative splicing plays an important role in key biological processes such as cell differentiation, lineage determination^2^ and tumorigenesis^3^, and its dysregulation has been connected to many diseases such as cancer^4^, intellectual disability and autism spectrum disorders (ASDs)^5^. While alternative splicing has been widely studied at the tissue level^2, 6, 7^, it remains poorly understood at the single cell level^8, 9^.

Technical and computational challenges have limited our ability to explore alternative splicing at single cell and spatial resolution. Commonly used droplet-based scRNA-seq protocols^10-13^ are based on short read sequencing that only captures the 3’ or 5’ ends of RNA transcripts. The Smart-seq protocol^14-16^ can achieve full-length transcript coverage but requires further assembly and has limited throughput. Advances in full length sequencing methods provide new opportunities for single cell isoform analysis, and recent studies have demonstrated their feasibility^17-20^. The two most commonly used full length sequencing protocols are provided by Pacific Biosciences (PacBio) and Oxford Nanopore Technologies (Nanopore). Current versions of Pacbio’s protocol have higher sequencing accuracy but limited sequencing capacity^21^, and thus cannot attain the coverage needed for transcriptome-wide single-cell or spatial alternative splicing analysis. In contrast, current Nanopore protocols have lower sequencing accuracy but can achieve higher sequencing capacity to profile a large number of transcripts^22^ across a large number of cells/spots. In this paper, we develop a computational pipeline for pre-processing and analysis of single cell and spatially barcoded Nanopore sequencing data. The data we analyze were generated by the 10x single cell and Visium platforms, but the model concepts should be generalizable to other platforms.

A key challenge in the analysis of single cell and spatially barcoded Nanopore sequencing data is the recovery of cell/spot barcode and unique molecular identifier (UMI) from each read under low per-base sequencing accuracy (87%-95%)^22-24^. These barcodes are short sequence tags which are not tolerant of sequencing errors. Proposed fixes to this problem include iterative sequencing^24^ to lower sequencing error, or modifying the read structure to increase error tolerance^23^. A third alternative is to sequence the same cDNA library with both short read and long read methods, and then use the short reads to construct reference barcode sets to help with the extraction of cell barcodes and UMIs from the long reads^19, 20^. While a shallow short-read sequencing run is sufficient for the construction of a reference cell/spot barcode list, the construction of a reference UMI list requires highly saturated sequencing, which can be prohibitive in cost. To avoid the need for saturated short-read sequencing, sicelore2^19^ and FLAMES^20^ cluster UMIs within a fixed edit distance to quantify isoform expression, but we will show that such a strategy is not foolproof: First, if sequencing error is high, existing methods overestimate the number of UMIs, causing inflated quantification. Such inflation due to sequencing error can be more severe with increased PCR amplification fold. Second, high gene expression causes “barcode crowding” in the UMI space, leading to erroneous merging of distinct UMIs and deflated quantification.

Low sequencing accuracy and read truncation can also lead to misidentification of isoforms. With the accumulation of sequencing errors in long reads, short exons are sometimes wrongly mapped, leading to the false positive discovery of novel isoforms. Read truncation, especially at the 5’ end due to pore-block and early stop in reverse transcription^25^, is also pervasive. These issues need to be addressed to allow reliable downstream alternative splicing analysis with single cell and spatially barcoded long read sequencing.

To address these myriad challenges, we developed an isoform-annotation-free computational pipeline to perform accurate and fast cell barcode extraction, UMI assignment, and mapping and truncation error correction. We validated this pipeline using simulated and real datasets, showing that it leads to accurate quantification of isoform expression within individual cells/spots. We then applied this pipeline to multiple single cell and Visium Nanopore sequencing datasets and developed a rigorous statistical framework to quantify splicing variation at the single cell/spot level. In particular, we decompose heterogeneity in the usage of an exon (or splice-site) into intra-cell/spot and inter-cell/spot splicing heterogeneity: For a given gene, intra-cell heterogeneity is the diversity of exon-usage within a cell, and inter-cell heterogeneity is the variation in exon-usage between cells. This framework gives a natural metric for the identification of exons with high inter-cell/spot splicing variation, which, similar to the pervasive concept of “highly variable genes”, can be used to profile splicing changes within the cell population or across a tissue section. On matched single cell and Visium Nanopore sequencing of the same tissue, that Longcell identifies highly variable isoforms that are in high agreement between the two modalities.

We further developed a new statistical procedure to the identification of exon-inclusion differences between cell populations, and applied it to a splicing factor knock-out experiment to identify novel targets for each splicing factor. The statistical procedure allows for rigorous type I error control and sensitive detection of changes in splicing distributions beyond a mean shift. The new methods and experimental data enable the characterization of regulation patterns of splicing factors at single cell resolution.

## Results

### Isoform quantification within single cells and spatial spots

For precise single-cell and spatial isoform quantification, we need to first recover the cell/spot barcode and the UMI sequence from the long reads. This task is complicated by the high sequencing errors rates, including the high prevalence of indels in Nanopore sequencing. Computational methods have been proposed to recover the cell barcode for Nanopore reads with high accuracy^19, 20, 26^ (Supplementary Fig 2D). However, there still a lack of accurate and efficient algorithm for UMI barcode recovery, necessary for reduction of PCR amplification noise, and truncation and mapping error correction, necessary for precise isoform quantification and alternative splicing detection. To illustrate how sequencing errors could influence the UMI sequence, we extracted the 10 base-pair sequence located 5’ of confidently assigned cell barcodes, which should be part of the Illumina R1 adapter (Fig 1A). Since the adapter sequence is known, we can compute the edit distance between the original adapter 10mer with the extracted 10mers. Although most extracted sequences have low edit distance from the original, it is notable that around 20% have edit distance greater than 2 (Fig 1A bottom left), which exceeds the threshold of existing barcode clustering procedures and causes bias in expression estimation. Using the observed sequencing error distribution as a guide (Fig 1A bottom right), we simulated UMI barcodes with comparable edit distances from the truth, across varying gene expression levels and PCR amplification folds. The first problem revealed by these simulations is that, in many cases, the observed UMIs corresponding to a single true UMI split into many small or singleton clusters due to the accumulation of sequencing errors (Fig 1C). This phenomenon, which we call UMI *scattering*, leads to inflated estimation of gene expression. The second problem is that the edit distance between true UMIs decreases with gene expression. This phenomenon, which we call UMI *crowding* (Fig 1B, C), makes it difficult to distinguish between reads corresponding to different molecules (different UMIs) versus reads coming from the same molecule but corrupted by sequencing errors (same UMI). For highly expressed genes, UMI crowding leads to the merging of clusters of observed UMI sequences, each corresponding to a distinct true UMI, and an underestimation of the true expression level (Fig 1C right).

**Figure 1:**
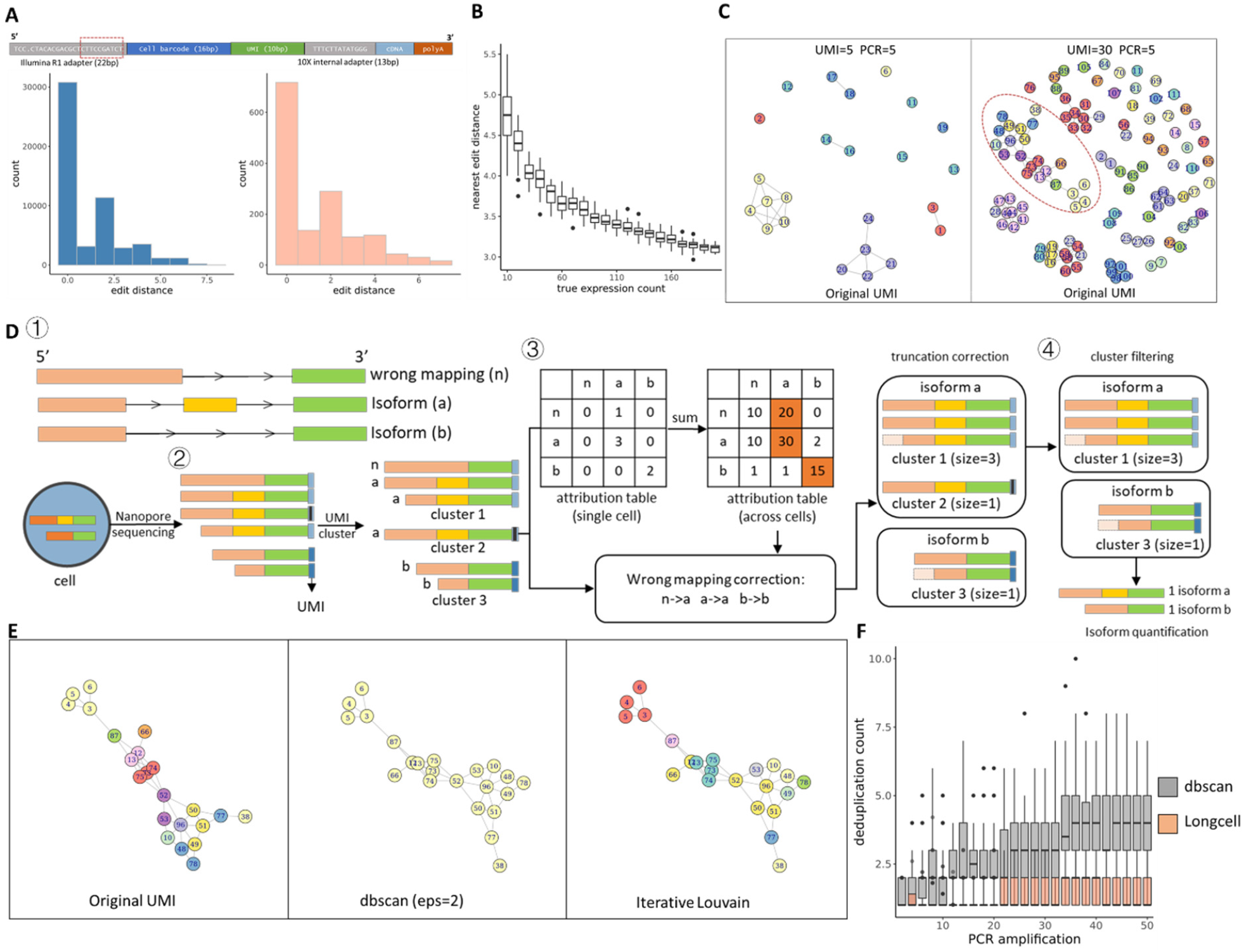
Overview of single cell Nanopore RNA seq preprocessing. **A** The structure of a read. 10mers aside the cell barcode (marked by red box) is extracted from sequences in real data. Then we compare those 10mers to their original sequences, and their edit distance distribution indicates the influence of sequencing errors to the UMI (bottom left). Bottom right shows the distribution of edit distance for the same region in our simulated sequences, which is very similar to that in real data. **B** Expected nearest edit distance between true UMI sequences of a gene at varying expression. 10 bp UMIs are randomly sampled at increasing counts to mimic the UMIs representing a single isoform in a single cell. As expected, as gene expression increases, the nearest neighbor edit distances between UMIs decrease. For example, since an insertion or a deletion can lead to an edit distance of 2, two different UMIs with edit distance lower than 4 can be connected by a “bridge” UMI sequence to which each differs by only an insertion or a deletion. The probability of such linkage increases with increasing gene expression. **C** UMI graph. Each node is a UMI after amplification with simulated sequencing errors to mimic the situation in Nanopore long reads. The color indicates if they are amplified from the same original UMI. UMIs with edit distance lower than 2 will be connected by an edge. Left: Simulation with 5 original UMIs and mean PCR amplification fold of 5. Different UMI clusters are separated. There also exist some singletons which is away from its original UMI cluster. Right: Simulation with 30 original UMIs and mean PCR amplification fold of 5. More singletons emerge in such high expression condition. Different UMI clusters are connected to each other by some bridge nodes into a connected sub graph (as marked by red circle). **D** UMI deduplication procedure: (1) simulated isoforms for a gene for illustration, including two types of isoforms (abbreviated as a and b) and a fake isoform due to wrong mapping (abbreviated as n). (2) UMIs are first clustered within each single cell. (3) Then isoform clusters are compared across all cells to correct for wrong mapping. Truncation errors are corrected by comparing to complete read within each cluster. (4) After the correction of truncation and mapping errors, small UMI clusters may be pruned based on the distribution of cluster sizes for clusters involving the same isoform. **E** We applied different clustering methods on this graph and show the clustering result on the marked sub graph in Fig 1.C right to see if they could separate the different UMI clusters. Left: Zoom in to the marked UMI sub graph, the color is the same as Fig 1.C right to indicate the original UMI. Middle: clustering result for dbscan method (eps=2) on the marked sub graph. All UMIs are clustered into one group. Right: clustering result for iterative louvain method on the marked sub graph. Most UMI clusters are recovered. **F** We sample 10mer adapters in different number (2∼50) to mimic PCR replicates of a UMI, then apply dbscan cluster (eps = 2, merge UMIs with edit distance lower than 2) and Longcell to sampled 10mer adapters to do deduplication. The dbscan method leads to inflated UMI estimation when amplification fold gets higher, while Longcell has a stable estimation even under the high amplification condition.

Addressing the above issues, we designed a new UMI recovery procedure which, simultaneously, utilizes PCR duplicates to correct truncation and mapping errors. As the UMI is located adjacent to the cell barcode, after cell barcode assignment we extract the adjacent sequence that putatively contains the UMI, together with short flanking sequences on both sides so as to account for possible insertions and deletions. Using the cell barcode, we first grouped all putative UMI sequences assigned to the same cell. Ideally, we should only need to disambiguate between UMIs mapping to the same isoform within the same cell, however, since isoform assignment is often misled by read truncation and wrong mapping (Supplementary Fig 1), the putative UMIs extracted for each cell are first grouped at a meta-isoform level. A meta-isoform group comprises of isoforms which could be transformed to each other through end truncations or wrong mapping of short middle exons. Within each meta-isoform group, we applied an iterative clustering procedure to cluster the putative UMI sequences (Fig 1D.2). The iterative clustering procedure is robust to UMI crowding (Fig 1E). Each cluster represents one unique original UMI, and thus correspond to one RNA molecule of origin.

**Figure 2:**
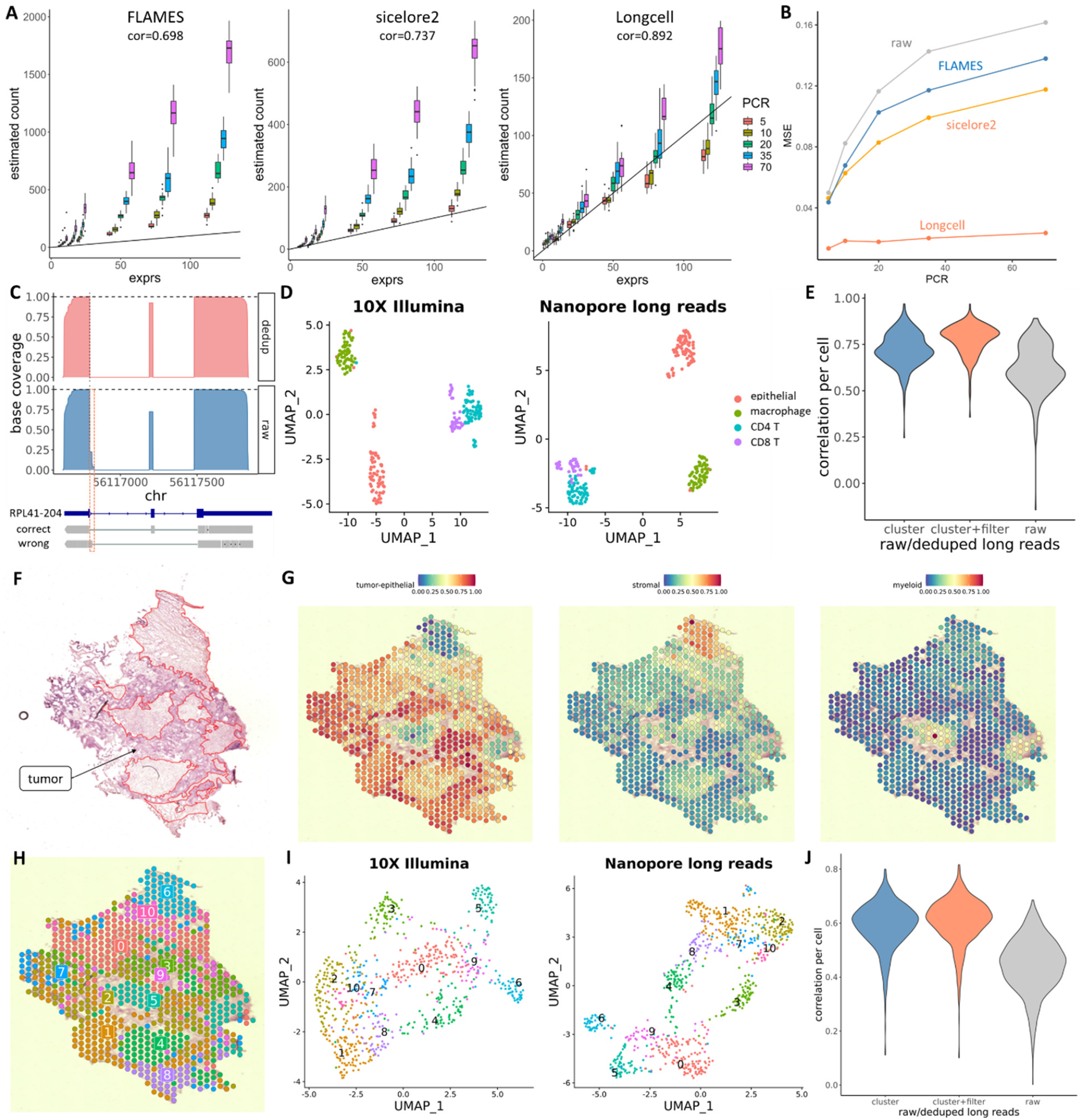
UMI deduplication results on simulated and real datasets. **A** The comparison of gene quantification by different methods on the simulated data across different amplification fold and gene expression. **B** The comparison of psi estimation by different methods on the simulated data across different PCR amplification fold, evaluated by mean square error. **C** Correction of wrongly mapped exons for RPL41-204 after UMI deduplication. **D** UMAP built on gene expression from 10X and paired ONT scRNA sequencing of the CRCLM sample. **E** Gene-wise correlation with Illumina-based estimates, per cell, for raw and processed ONT-based estimates. **F** The histological annotation for the CRCLM sample. Non-tumor regions are marked with red circles. **G** Cell type composition in each Visium spot (Left: tumor epithelial. Middle: stromal. Right: Myeloid). **H** The spatial view of the Visium spot clustering. **I** UMAP built on gene expression from long reads and paired short reads. **J** The per spot correlation between long reads Visium sequencing and paired short reads Visium sequencing for a CRCLM sample.

After clustering, the longest dominant isoform within each meta-isoform group is taken as the representative isoform for that cluster. This step corrects most truncation and mapping errors, as wrongly mapped/truncated reads are usually in the minority within each UMI cluster (Fig 1D.3). Finally, we hypothesize that all expressed transcripts of the same isoform should have similar amplification fold, as they share the same RNA sequence (Supplementary Fig 3A), and therefore, they should have similar cluster sizes. Thus, within the same isoform, we rank clusters by size and clusters that are too small (e.g. singletons) are removed as they are likely created by sequencing errors. We found that this filtering step addresses the UMI scattering issue and substantially improves expression estimation (Simulation results in Fig 1F, validation in real single cell and Visium data in Figure 2E and J).

**Figure 3:**
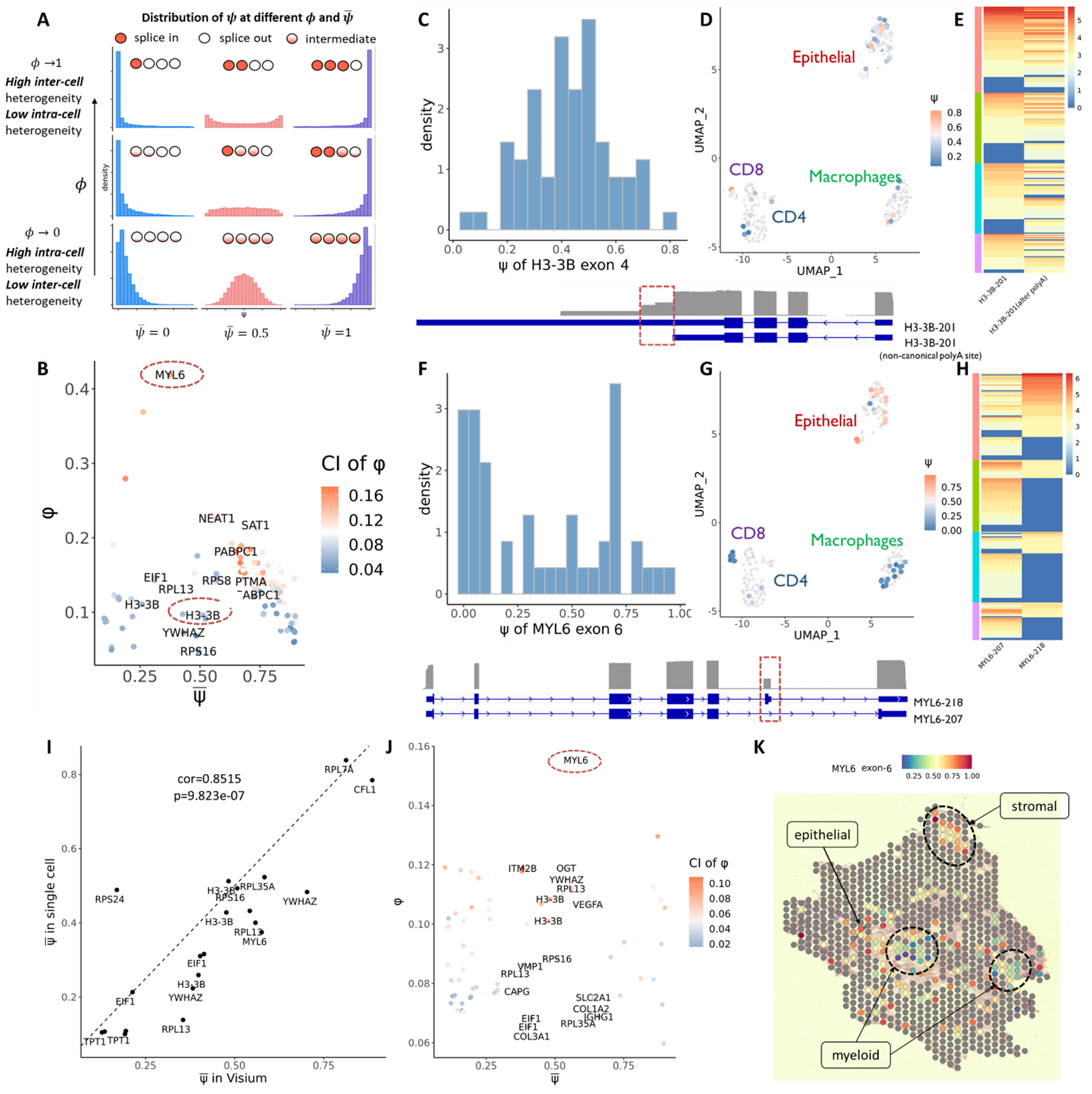
Quantification of intra-cell versus inter-cell isoform heterogeneity in colorectal metastasis to the liver. **A** The relationship between 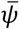 and *ϕ*, 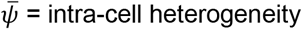 intra-cell heterogeneity, *ϕ*= inter-cell heterogeneity.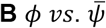 distribution for alternative spliced exons in CRCLM single cell data, the color indicates the confidence interval of *ϕ*. Circles indicate two examples with low and high *ϕ*. **C** The *ψ* distribution for exon 4 of H3-3B, which has a relatively low *ϕ*, indicating low intercell heterogeneity. **D** The *ψ* distribution for exon 4 of H3-3B across cells, showing similar inclusion-level of this exon across all cells. **E** Distribution of the two dominantly expressed isoforms across different cell types, both isoforms are co-expressed in each cell at a similar ratio. **F** The *ψ* distribution for exon 6 of MYL6, which has a very high *ϕ*, indicating high intercell heterogeneity. **G** The *ψ* distribution for exon 6 of MYL6 across cells, epithelial shows the highest inclusion-level of this exon. **H** Distribution of the two dominantly expressed isoforms across different cell types, epithelial has higher expression of MYL6-218 compared to other cell types. **I** The correspondence of 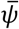 between single cell long reads and Visium long reads sequencing. **J** The relationship between 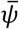 and *ϕ* in CRCLM Visium sequencing. **K** The spatial view of *ψ* for exon 6 of MYL6, the aggregation regions of myeloid are marked by blue, which show low inclusion of this exon, while the aggregation region of stromal is marked by red circle. The rest is tumor epithelial. Both epithelial and stromal region show high inclusion of exon 6 of MYL6.

We evaluated whether our UMI deduplication method gives accurate isoform quantification under different PCR amplification folds and sequencing error rates, in simulated data with the same edit distances as in Fig 1A. Compared to sicelore2^19^ and FLAMES^20^, Longcell can remove the amplification bias more effectively, with the estimated gene expression mostly keeping a linear relationship with real gene expression across different amplification folds (Fig 2A). Compared to existing state-of-the-art, Longcell also provides a more accurate exon percent-spliced-in (psi) estimation after deduplication (Fig 2B).

As proof-of-principle, we first analyze a sample of colorectal cancer metastasis to the liver (CRCLM) which was processed by both 10x Chromium (single cell) and 10x Visium (spatial), with the cDNA libraries from both samples submitted to Oxford Nanopore and Illumina sequencing. The matched 10X Illumina sequencing data for this sample was used as a benchmark in both the single cell and Visium samples. We first prove that Longcell is able to correct read mapping and truncation errors without relying on existing isoform annotations. An example is given for the RPL41 gene, where a small middle exon that was wrongly mapped in 25% of the reads was corrected by Longcell (Fig 2C), recovering the existing isoform annotation for this gene without a priori knowledge of it. Then for the gene quantification in single cell sequencing, the Umaps derived from Illumina short-read gene expression and Nanopore long-read gene expression reveal similar geometries (Fig 2D), indicating that the cell barcode extraction and UMI recovery from the Nanopore sequencing data successfully recovered the underlying biological variation from the background noise. The per cell correlation of long- and short-read based gene expression estimates given by Longcell was significantly higher than the correlation given by raw read counts (Fig 2E); in particular, the algorithm for reducing UMI scattering significantly improves correlation with short reads at the gene expression level, indicating that this step is important for quantification accuracy.

Now consider the Visium sample. Fig 2F shows the histology of this tissue section, with expert pathology annotation of the tumor regions. A cell-type deconvolution of the spots reveals the regions with high tumor cell concentration, agreeing with the pathology annotation. The deconvolution also high-light the regions of high myeloid and stromal cell content. Fig 2I shows, respectively, the Umaps of the short and long read data sets generated from this section, colored by a joint clustering of these two data modalities. We see that, as the single cell data set, the geometry of the two modalities agree, with clusters in the Illumina short read data also well separated in the Nanopore long read data. These clusters map to contiguous regions in space (Fig 2H), adhering to the boundaries of the tumor, stromal, and myeloid regions delineated by the pathology annotation (Fig 2F) and the in silico deconvolution (Fig 2G). This proves that Longcell is able to extract meaningful biological structure from long read sequencing of this Visium cDNA library. Finally, a comparison at the gene-level between the short and long read data sets reveals that Longcell improves expression quantification, as compared to using library-size normalized counts. As for the single cell data set, significant improvement in quantification accuracy is achieved by the scattering-reduction algorithm.

### Quantification of intra and inter-cell splicing variation in spatial and single cell data

Based on the Longcell preprocessing procedure, we can now characterize the isoform-level heterogeneity across cells and spatial positions within a tissue. When an alternative splicing event is detected at the bulk-tissue level, a basic question is whether the two isoforms are co-expressed within the same cell or in separate cells. This question has been examined by single cell short-read sequencing^27, 28^, where the conclusion was that alternative isoforms predominantly originate from different cells. However, these short-read based studies were plagued by low sequencing coverage^29^ that limits their power to detect alternative splicing within the same cell. Here, we propose a statistical model to explicitly quantify inter-cell/spot splicing variation at a given exon/splice site, and to measure the degree of inter-versus intra-cell/spot splicing heterogeneity: *Intra*-cell/spot heterogeneity refers to the case where alternative isoforms co-exist within the same cell/spot, whereas inter-cell/spot heterogeneity refers to the case where the splicing profile of a gene differs across cells/spots.

Consider a given exon within a given gene. For cell *c*, let *X*_*c*_ be the count of reads with this exon spliced in. We assume that *X*_*c*_ ∼ *Binomial*(*N*_*c*_, *ψ*_*c*_), where *N*_*c*_ is the total number of reads mapping to the gene in the given cell, and *ψ*_*c*_ is the cell-specific “percent-spliced-in” parameter, i.e. the proportion of transcripts of cell *c* with the exon retained. With *ψ*_*c*_ unobserved, our goal is to characterize the distribution of *ψ*_*c*_ across cells. Since *ψ*_*c*_ ∈ [0,1], we use the Beta distribution to flexibly model its variation across cells. The mean of the Beta distribution, *μ = E*[*ψ*], is the bulk-level percent-spliced-in for this exon, while the variance of the Beta distribution measures the amount of inter-cell variation in exon usage. Since the variance of the Beta distribution scales with *μ*(1 *– μ*), we compute the dispersion *ϕ = var*(*ψ*)[*μ*(1 *– μ*)]^−1^ as a mean-invariant measure of inter-cell splicing heterogeneity. *ϕ* is a continuous value with range [0,1]. A low *ϕ* indicates that the distribution of *ψ* across cells is unimodal, and thus individual cells express both isoforms at a similar ratio across cells. In contrast, a high *ϕ* indicates that *ψ* is bimodal, meaning that individual cells splice this exon in or out in a binary way (Fig 3A). By Bootstrap, we can compute the standard error for 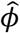. For a given dataset, a plot of *ψ* versus *ϕ* across genes and exons gives a birds-eye-view of mean exon inclusion rates versus the degree of inter-cell/spot splicing variation across the transcriptome. This method of estimating inter-cell splicing variation adjusts for differences in per-cell coverage across sites. See Methods for model details and estimation and inference procedure.

The above model was fit to highly expressed genes in the CRCLM dataset. The standard error in *ϕ* is shown as continuous color scale in the *μ* versus *ϕ* plot. Contrary to published literature, we found that most alternatively spliced exons for highly expressed genes have low estimated *ϕ* indicating a low inter-cell heterogeneity but high intra-cell heterogeneity (Fig 3B). Thus, for highly expressed genes, the dominant pattern is for different isoforms to be co-expressed at a similar ratio within each cell across the cell population. For example, we observed a non-canonical alternative poly(A) site in H3-3B-201 (Fig 3C). This isoform has a long exon at its 3’ end, but nearly half of the transcripts show early stop at the middle of this exon in each cell, with little inter-cell heterogeneity. Such high intra-cell heterogeneity was also observed in other samples, through both full transcriptome and targeted gene sequencing (Fig 4A and Supplementary Fig 4).

**Figure 4:**
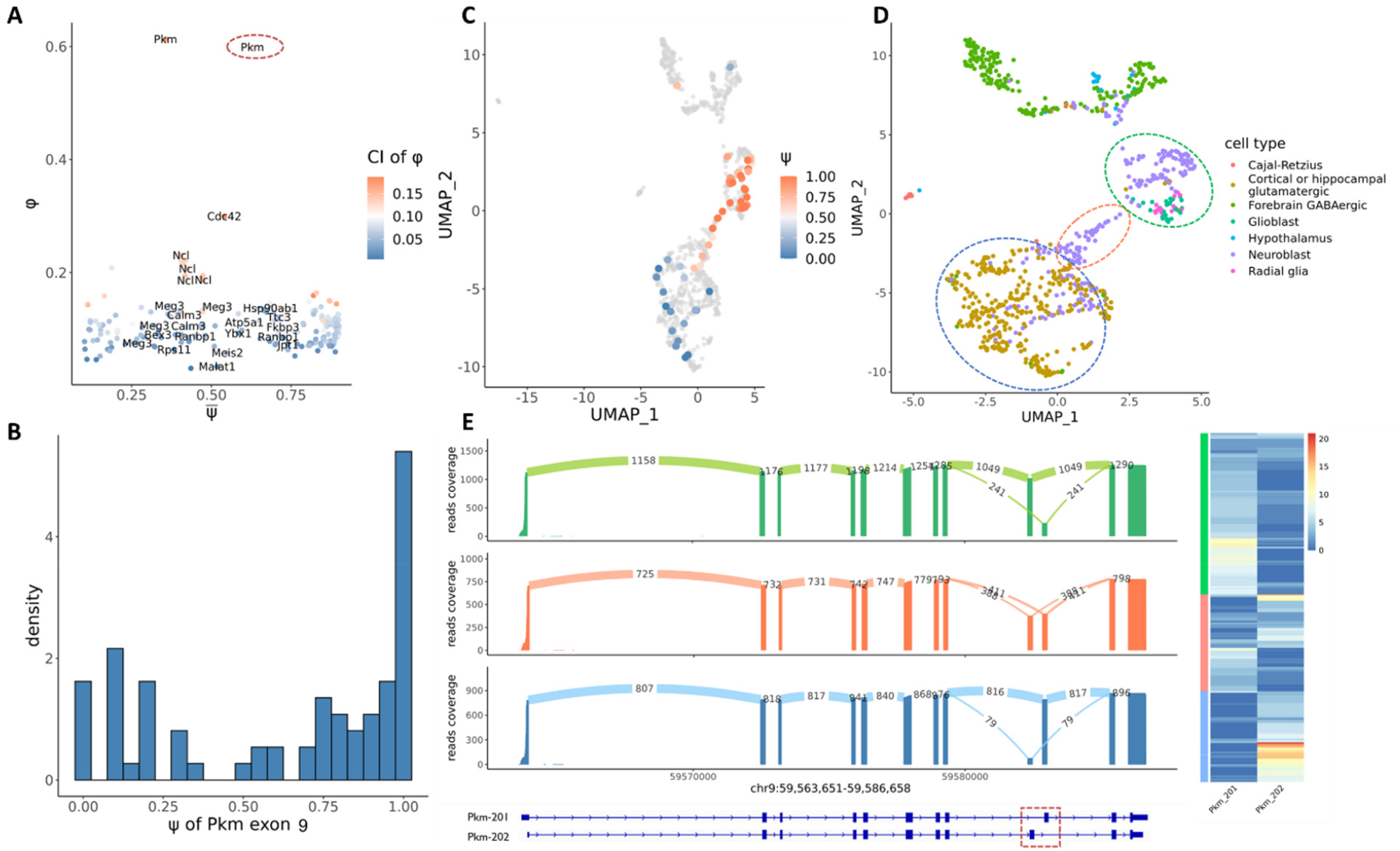
Quantification of intra-cell versus inter-cell isoform heterogeneity in embryo mouse brain. **A** *ϕ vs*. 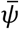 distribution for alternative spliced exons, color indicates the confidence interval of *ϕ*. The circle indicates the high *ϕ* example Pkm exon 9. **B** The *ψ* distribution for exon 9 of Pkm, which has a relatively high *ϕ* and show a bimodal distribution, indicating a high inter-cell heterogeneity. **C** Umap of cells in mouse embryo brain. Cells are colored by *ψ* for exon 9 of Pkm. Cells which have low expression (<10) of this gene and couldn’t give a confident *ψ* estimation are colored in grey. **D** Umap of cells in mouse embryo brain. Cells are colored by cell types. Three parts of cells are circled according to their *ψ* for exon 9 of Pkm. **E** The alternative splicing for Pkm in above three circled groups. The alternative splicing of exon 9 in Pkm mainly leads to 2 isoforms: Pkm-201 ad Pkm-202. An obvious transition of the expression of two isoforms can be identified both in both bulk (sashimi plot at left) and single cell level (heatmap at right).

**Figure 4:**
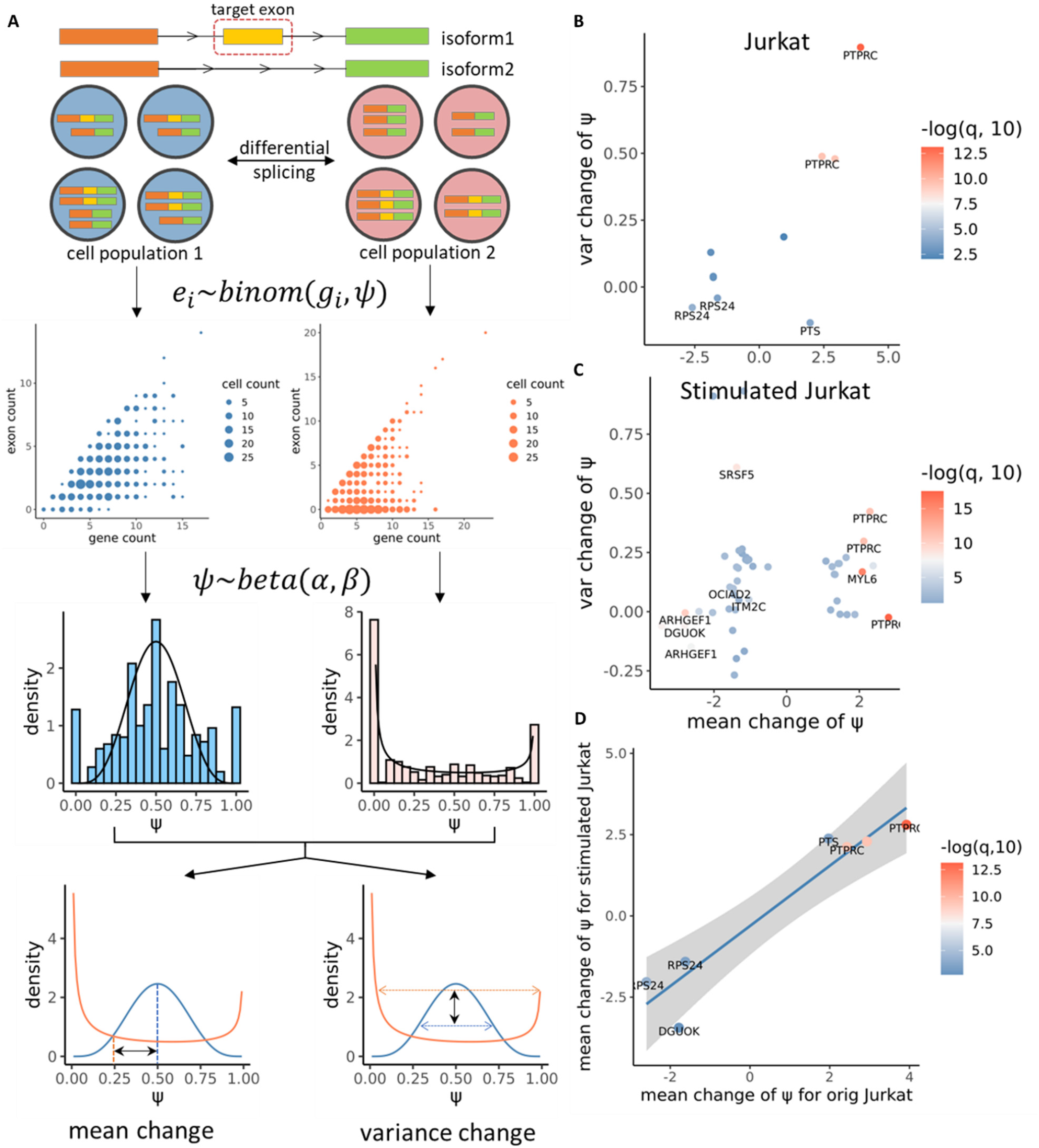
Differential splicing analysis and the detection of alternative splicing regulated by splicing factors. **A** The principle of generalized likelihood ratio test to identify differentially expressed isoforms. **B**,**C** All significant exons identified in Jurkat (B) and stimulated Jurkat cells (C) after knock-out of spicing factors. **D** Correspondence of significant exons between original and stimulated Jurkat cells. After stimulation there is a significant change of gene expression and alternative splicing in Jurkat cells, but the regulation of splicing factors keeps the same direction.

Meanwhile, we found alternatively spliced exons with high *ϕ* indicating high inter-cell variation and a bimodal distribution for *ψ*. An example is exon 6 in MYL6. The two isoforms of MYL6 are enriched in different cell groups, with dominance of MYL6-218 in tumor epithelial cells and dominance of MYL6-207 in macrophages and CD8 cells (Fig 3F, G and H). Just as highly variable genes analysis identifies genes with high variation in total expression across cells, inter-cell splicing variation analysis identifies exons with high variation in percent-splice-in across cells and is useful in revealing differentially expressed isoforms across the cell population.

For the same CRCLM sample analyzed above, we performed Visium spatial transcriptome profiling with spot-matched Illumina and Nanopore sequencing. As the single cell data, correlation between the short and long read estimates of spot gene expression significantly improved with Longcell preprocessing procedure (Supplementary Fig 3B). UMAP embedding of spots from the VISIUM sample sequenced by Illumina and Nanopore display similar geometries, with cell clusters identified in the Illumina modality even better separated in Nanopore modality (Supplmentary Fig 3C, D). This validates that Longcell effectively separates biological signal from noise in Visium barcoded Nanopore sequencing data. Spatial plot of the VISIUM sample shows that the Nanopore-derived clusters coincide with deconvolution-based labeling of the spots into tumor (epithelial), myeloid and stromal regions.

As for the single cell data, we estimated the cross-cell mean (*μ*) and dispersion (*ϕ*) for exon- and cell-specific *ψ* in highly expressed genes in the Visium Nanopore data. The mean *μ* in Visium highly correlates with those in single cell data (Fig 3J). As for the dispersion, in this case of Visium, a high value of 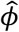 reflects high inter-spot heterogeneity in the splicing of an exon. Genes with a high dispersion in the single cell data, such as MYL6, also has high dispersion in the Visium data (Fig 3K), although overall, *ϕ* is lower in the Visium data due to cell aggregation within each spot. In the spatial view of the *ψ* distribution for MYL6 exon 6, we found that regions enriched for tumor cells show very high inclusion of this exon, while regions enriched for myeloid cells show low inclusion rates (Fig 3L). This is concordant with the cell-type specific splicing patterns for MYL6 detected in the matched single cell data, and thus, the results from the two modalities validate each other.

We further applied Longcell to Nanopore sequencing data of mouse embryonic brain cells^19^. As in the CRCLM data, high intra-cell and low inter-cell splicing heterogeneity was also observed in this dataset for most of the alternatively spliced exons (Fig 4A). Contrary to most alternatively spliced genes, we found the Pkm gene to display very high inter-cell splicing heterogeneity (Fig4 B), which corresponds to the published result that single cells tend to dominantly express one transcript isoform for the Pkm gene in this tissue^19^. Our analysis found that exon 9 for Pkm has a continuously increasing trend towards retention as cells progress from neuroblasts to glutamatergic neurons (Fig4 C, D). This indicates a continuous switch in splicing from the Pkm-201 isoform to the Pkm-202 isoform (Fig4 E) as neurons mature towards the differentiated state.

### Detection of differential splicing events between cell populations

We next developed a method for the identification of differentially spliced genes between two cell populations. Consider the exon colored in yellow in Figure 4A. As in the last section, we model its count *X*_*c*_ for each cell *c* with a Beta-Binomial distribution, *X*_*c*_ ∼ *Binomial*(*N*_*c*_, *ψ*_*c*_), with *ψ*_*c*_ ∼ *Beta*(*α*_1_, *β*_1_) for cells in population 1, and *ψ*_*c*_ ∼ *Beta*(*α*_2_, *β*_2_) for cells in population 2. Based on this model, we perform a generalized likelihood ratio test of the null hypothesis *H*_*O*_: *α*_1_ *= α*_2_, *β*_1_ *= β*_2_ which corresponds to the scenario where the two cell populations have the same Beta distribution parameters for the cell-specific percent-spliced-in values. This model can account for uneven total gene expression across cells and, by testing for changes in both shape parameters of the underlying Beta distribution, can sensitively detect both a shift in mean and a change in dispersion in the population distribution of *ψ*_*c*_. After correction for multiple hypothesis testing, exons that display differences in the distribution of *ψ* across the two cell populations are reported. The detected signals will then be decomposed into mean-level and variance-level changes for better interpretability (Fig 4A).

### Longcell reveals targets of splicing regulators in single cell CRISPR experiment

We applied Longcell, with the differential expression module, to a perturbation experiment in naïve and stimulated Jurkat human cell lines. The Jurkat cell line was derived from a T cell leukemia and stably expressed Cas9. On this cell line, we transduced a multiplexed gRNA lentiviral library targeting 9 splicing factor genes (two gRNAs per gene), including well-known factors from the HNRNPLL and SRSF families and less-studied factors such as PCBP2 and CWC27 (Supplementary Fig 5B). After 14 days, we harvested the cells, generated single-cell libraries, and conducted full transcriptome Nanopore sequencing (Supplementary Fig 5A) ^30^. We identified 10 exons in 5 genes in Jurkat cell lines and 50 exons in 20 genes in stimulated Jurkat cell lines that were affected by the knock-out of splicing factors after FDR control at threshold 0.05 (Fig 4B,C and Supplementary table 1). For example, we observed that knock-out of HNRPLL significantly promoted the inclusion of exon 4 in PTPRC, while knock-out of CELF2 promoted the inclusion of exon 6 in MYL6, which corroborates existing knowledge about these factors^30, 31^ (Supplementary Figure 6).

**Figure 5:**
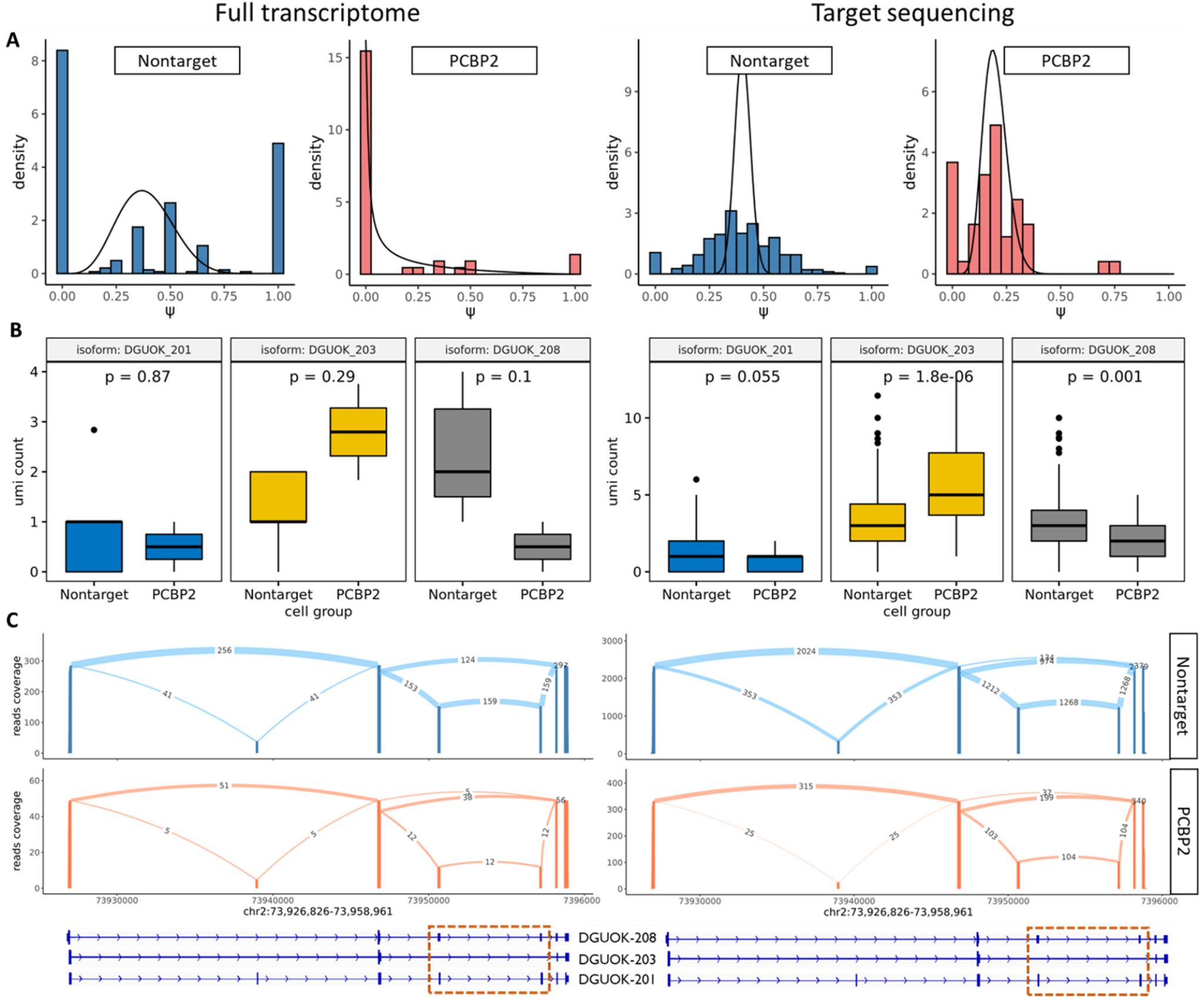
Isoform transition of DGUOK after knock-out of PCBP2. **A** *ψ* distribution of exon 3 and 4 of DGUOK in nontarget and PCBP2 knock-out cells. An obvious decreasing of *ψ* can be observed after knock-out. Left for the full transcriptome and right for the target sequencing. **B** Comparison of expression for 3 main DGUOK isoforms in nontarget and PCBP2 knock-out cell populations. Both full transcriptome (left) and target sequencing (right) show the same pattern. **C** Sashimi plot for 3 main DGUOK isoforms in Nontarget and PCBP2 knock-out cell populations.

We also compared the regulation patterns of splicing factors between stimulated and unstimulated T cells. Although the stimulation significantly changed the expression profile of T cells, we were still able to identify overlapping regulation signals between the naïve and perturbed settings. For these overlapping targets, the effect sizes of splicing factor knockout correlate highly between the stimulated and unstimulated group (*p =* 0.001), validating our procedure and also revealing that these specific regulatory relationships of the splicing factors are not changed by the stimulation (Fig 4D).

Besides the well-known targets mentioned above, we also identified novel regulation patterns such as PCBP2 promoting the inclusion of exons 3 and 4 in DGUOK (Fig 5A). Knock-out of this splicing factor can lead to the transition from the expression of isoform DGUOK-203 to isoform DGUOK-208 (Fig 5B, C). Due to the limited depth of the DGUOK gene in the full transcriptome, this signal is weak. To validate this finding, we performed targeted sequencing of DGUOK from the same cDNA library. We applied Longcell to the targeted sequencing data to quantify the expression of DGUOK-203 and DGUOK-208 in the non-target and PCBP2 knock-out cells, and identified significant splicing change from DGUOK-203 to DUOK-208 after knock-out of PCBP2 (Fig 6B right). This result also confirms that Longcell can sensitively detect subtle changes in splicing distribution between cell populations.

## Discussion

Long read sequencing is a powerful tool for detecting and characterizing alternative splicing in single cell and spatial transcriptomics. However, the influence of sequencing, read truncation, and mapping errors in high throughput long-read data on the identification and quantification of isoforms has not been well characterized. Here, we performed detailed study of the influence of these sources of noise, and described two phenomena, *UMI crowding* and *UMI scattering*, which can severely bias isoform quantification. We also observed that wrong mapping and technical truncation of exons can lead to spurious alternative splicing detections. To overcome these issues, we developed Longcell, a novel preprocessing method that efficiently provides accurate isoform quantification for long-read sequencing with single-cell and spatial barcodes. Longcell is benchmarked in simulations under a broad set of settings. Longcell is also validated by comparing cell-matched Illumina- and Nanopore-based estimates, by comparing sample-matched Visium and single cell Nanopore sequencing results, and by targeted Nanopore sequencing of select alternative splicing signals.

Our results inform experimental design for future single cell long-read studies: Although Longcell partially address the UMI crowding issue, complete removal of crowding-related biases would require the design of sequencing protocols utilizing longer UMI barcodes. Also, it is important to note that we correct for read truncation and mapping errors by taking a “consensus” among reads assigned to the same UMI. PCR replicates are both a blessing and a curse, and our simulations show that a PCR amplification fold of 5 is enough to correct wrong mapping. Thus, a shallow PCR amplification before sequencing is still recommended.

For downstream analyses, we proposed a novel metric to evaluate the splicing heterogeneity in cell populations. Armed with this metric, we re-examined the question of intra-cell splicing heterogeneity that was the focus of previous studies^27-29^: Do cells predominantly express only one isoform, or do they co-express multiple isoforms of the same gene? Across multiple datasets, we found that co-expression of multiple isoforms is the dominant pattern across highly expressed genes. Many of these alternative splicing events involve alternative 3’ polyadenylation. Further research is required to reveal the biological mechanisms of intra-cell splicing heterogeneity.

We also developed a new method for differential splicing analysis that detects, for an exon or splice-site, a change in the distribution of its cell-specific percent-spliced-in between two groups of cells. The test adjusts for differences in read coverage between cells, and goes beyond shifts in mean to detect changes in the shape of the distribution. We performed a splicing factor knock-out experiment, and applied Longcell to identify targets for multiple splicing factors, overcoming the severe sparsity of the full-transcriptome sequencing data.

In general, Longread overcomes severe challenges in cell barcode and UMI recovery and allows downstream analysis via a simple but effective model for splicing variation at single cell resolution. Through new experiments and re-analysis of existing data sets, we have demonstrated the feasibility of full-transcriptome profiling of alternative splicing patterns in single cells.

## Methods

### Wet experiment

#### Cell culture and CRISPR knock-out

Jurkat (ATCC TIB-152) cells were maintained in Roswell Park Memorial Institute (RPMI) 1640 medium supplemented with 10% FBS. Pooled splicing factors knock-out were performed as previously described^30^.

#### Single-cell library preparation and sequencing

Single-cell cDNA and gene expression libraries are generated by Chromium Next GEM Single Cell 5’ Library & Gel Bead Kit v2 (10X Genomics, Pleasanton, CA, USA) as per the manufacturer’s protocol. The sgRNA direct capture was performed as previously described^30^. We amplified cDNA and gene expression libraries with 16 and 14 cycles of PCR respectively. The quality of libraries was confirmed using 2% E-Gel (ThermoFisher Scientific, Waltham, MA, USA). They were then quantified by Qubit (Invitrogen). Gene expression libraries were sequenced using Illumina platforms (Illumina, San Diego, CA, USA). cDNA libraries were sequenced by Oxford Nanopore platform using Promethion flow cell FLO-PRO002 (Oxford Nanopore Technologies) as per the manufacturer’s protocol.

#### Nanopore reads preprocessing and mapping

We performed basecalling on the raw fast5 data using Guppy5. Nanopore reads were aligned to the human Genome (GRCh38) with minimap2 v2.24 in spliced alignment mode (command: “minimap2 -ax splice -k14 -t $thread –junc-bed $bed –secondary=no –sam-hit-only $refer $fastq > $output”). The splice junction bed file was generated from the Gencode v39 GTF using paftools.js, a companion script of minimap2.

#### CellBC and UMI assignment to Nanopore reads

1. Softclip extraction: After reads alignment, the 5’ and 3’ softclips will be extracted. For 5’ toolkit, the 55 bp segment proximal to the transcript within softclips will be preserved for downstream cell barcode and UMI alignment. The 3’ softclip will be used to detect if polyA exists (over 15 A or T must be found in an 20bp window). For 3’ toolkit, the 55bp segment after the polyA will be preserved. The start position for each softclip needs to be from the end that is proximal to the transcript.
2. Read filtering: Each read is recorded as a set of splice sites and its start and end position. Splice sites are counted across all reads for a gene. Reads with the splicing site whose count are lower than 10 will be filtered out.
3. Cell barcode match (Supplementary Fig 2A): The reference barcodes can be obtained from a matched Illumina sequencing of the cDNA library, or, in the absence of this Illumina run, one can use the 10X barcode pool that contains all possible barcodes. All barcodes will be vectorized into 8-mers to build an 8-mer dictionary. At the first iteration of barcode alignment, a prior distribution for barcode start position will be set as a normal distribution which could cover the whole softclip. The 95% confidence interval for this distribution should indicate the search region. Barcodes with 8-mer overlap with the search region will be preserved as candidates and will be further compared by edit distance. The barcode with the minimum edit-distance will be aligned and its start position will be used to update the start position distribution. After all softclips get aligned, alignment with high edit distance and deviant start positions will be filtered out as low quality alignment.
4. UMI clustering (Figure 1D, step 2): As UMI is known to be located beside the cell barcode, after cell barcode alignment, we extract this region that putatively contains the UMI, with 1bp flanking bases to be tolerant of insertions and deletions. We then define a meta-isoform group as the set of reads representing isoforms which could transform to each other by end truncations and wrong mapping of small middle exons (shorter than 80 bp). For example, for two reads *R*_1_, *R*_*2*_ with overlapping regions, here we define *s* as the start point of the read, and *e* as the end point of the read, *base*(*R*_*i*_, *a, b*) as the bases for read *R*_*i*_ within point *a* and *b*, then the overlapping region for the two reads should be *s =* max(*s*_1_, *s*_*2*_), *e =* min(*e*_1_, *e*_2_). *R*_1_, *R*_2_ belong to the same meta-isoform group if:

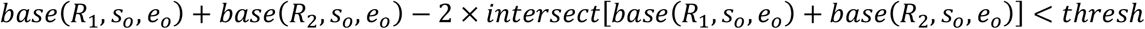

For each meta-isoform group within each cell, we calculate Needleman-Wunsch scores between UMIs to serve as similarity. We then build a graph based on the Needleman-Wunsch similarity. Each node is a UMI and the edge weight is the Needleman-Wunsch score. For 10bp UMIs, edges with weight lower than 5 will be filtered out. We then apply iterative Louvain algorithm on this graph using the cluster_louvain(graph, resolution = 1) function in R package igraph. Each time UMIs within the same cluster will be separated as a sub-graph. The iteration continues until all subgraphs are highly connected (min_cut > 3) or the minimum edge weight for all sub-graphs is larger than 7.
5. Wrong mapping correction (Figure 1D, step 3): As wrong mapping is mainly due to sequencing errors that occur after PCR amplification, wrongly mapped reads could coexist with correctly mapped reads in the same UMI cluster. Within each UMI cluster, the dominant and longest isoform is chosen as the representative for this cluster. An attribution table will be built to count how many reads for each isoform are assigned to this representative in this cluster. Such attribution table will be summed across all UMI clusters across all cells as an integration. As many wrong mapped reads coexist with the correct ones in the same UMI cluster and correct mapped reads should have quantity advantage, wrong mapped reads should be mostly assigned to the correctly mapped reads.
6. Scatter reduction (Figure 1D, step 4): Molecules of the same isoform should have similar PCR amplification fold, as they share the same oligonucleotide sequence (Supplementary Fig 3A). Thus, small clusters (e.g. singletons) that deviate from the “median” cluster size are likely to be due to the UMI scattering effect caused by sequencing errors. We use a statistical procedure to detect scattering, based on a negative binomial model for the UMI cluster size. To be more specific, suppose there are *n* transcripts for an isoform in a cell, then the UMI cluster size for each transcript follows a negative binomial distribution: (*C*_*i*_ *-* 1)∼*NegaBinomial*(*r, p*) according to our model (As the minimum of *C*_*i*_ is 1). There should be a linear relationship between its mean 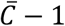 and variance *σ*^2^(*C*) as 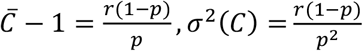. So here We have 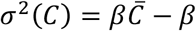. To estimate *β*, we performed the regression of *σ*^2^(*C*) versus 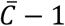, across UMI clusters for each gene in each cell, on Illumina short read data where we are confident of the UMI extraction. We limit the regression on data from genes with only one isoform, so that even with Illumina data we can be certain that all reads come from a molecule with the same isoform sequence. We constructed a 90% confidence interval for *σ*^2^(*C*) by asymptotic approximation (Supplementary Fig. 3):

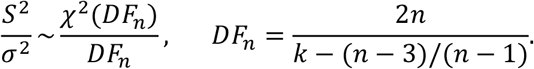

By the logic that the Nanopore data should have the same PCR amplification fold distribution as the Illumina data, we trim the small UMI clusters until the 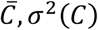, *σ*^2^(*C*) for the Nanopore data matches the estimated relationship above. That is, we iteratively remove the smallest UMI cluster until the confidence interval of the variance of UMI cluster size cover 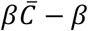.

### Single-cell gene expression quantification and cell type identification. Quantification of single cell isoform heterogeneity

1. Sub-exon definition: we cut exons for a gene into non-overlapping sub-exons according to the Gencode(V39) transcript annotation. Each corrected read after UMI deduplication is aligned to those sub-exons. When we assign each read to the sub-exons, the start and end site of the read can be allowed to be different from annotation as there are frequent truncations, but at over 10 bp overlap between the read and the sub-exons in the annotation are required. While middle splice sites are required to be no farther than 1 bp than the annotated sites.
2. Sub-exon count: For each read we use 0 to represent a sub-exon splice-out event and 1 for a spice-in. Then a read can be represented as a vector of 0 and 1 with the length as number of sub-exons to indicate its incorporation. As truncations at the 5’ are frequent in Nanopore long reads sequencing, it’s hard to figure out if a read is a new isoform with new transcription start site or it’s truncated. For robustness, we treat each read as censored data. Lack of exon at 5’ or 3’ without polyA won’t be considered as a splice-out event and will be record as NA. Thus, for each sub-exon *i* in a gene in each cell *c*, the gene count *G*_*ci*_for this sub-exon should be the count of non-NA values. While the exon count *E*_*ci*_ should be the count of 1.
3. Exon merge: Some exons are always coexisting or mutually exclusive, which contain the same information. Thus, individual analysis for those exons one by one can be waste of computation and can raise multiple testing problem. To merge those exons, we first identify coexisting and mutually exclusive exon pairs for each gene across all cells. If the coexist or mutual exclusiveness of those exon pairs can also be proved in the Gencode(V39) transcript annotation, they will be merged as a meta-exon.
4. Exon phi estimation: For accurate estimation of exon psi ad phi, we only pick meta-exons with *G*_*ci*_ ≥ 10 over in over 30 cell and estimate the psi value as 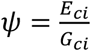. We then model exon psi with beta distribution *ψ*∼*Beta*(*α, β*) and use 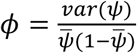 to control the correlation between mean with variance. To characterize the variability of the phi estimation, we use bootstrap to calculate the 95% confidence interval for each candidate meta-exon. S with 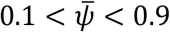 and *CI*(*ϕ*) <0.2 are preserved for downstream analysis.

#### Differential alternative splicing analysis

Here we model the distribution of the exon count for exon *i* for each cell *c* with a beta-binomial distribution: *E*_*ci*_∼*betabinom*(*G*_*ci*_, *α, β*). Based on this distribution, we could do a generalized likelihood ratio test to identify if two groups (*g*_1_ *and g*_2_) share the same parameters of *α* and *β*. The distance between the *ψ* distribution from two groups is quantified as Wasserstein distance. The confidence interval for the distribution distance is calculated by bootstrap. FDR control is applied to significant exons and only exons with *q* < 0.05 are preserved for downstream analysis.

